# Characterizing risky play-like behaviour in juvenile versus young adult rats

**DOI:** 10.64898/2026.10.29.757313

**Authors:** A. Souiki, J.S.D. Getty, S. Maggioncaldasacerdote, G.P. Lazzarino, F.J. Meye, E.J.M. Achterberg, H.M.B. Lesscher

## Abstract

Play is an evolutionarily conserved behaviour that contributes to healthy behavioural and neural development. While animal play is traditionally categorized as locomotor, object, or social play, developmental research in children has highlighted the importance of risky play, defined as voluntary engagement with uncertainty and manageable physical risk. Whether similar forms of behaviour occur in non-human animals remains largely unexplored. The present study aimed to establish an ethological framework for identifying and quantifying risky play-like behaviour in rats. Juvenile and young adult male Lister Hooded rats were repeatedly exposed to a custom-designed environment containing elevated and unstable structures that provided opportunities for climbing, balancing, hanging, jumping and social interaction. Behaviour was quantified using two complementary ethograms: a location-based ethogram describing the animal’s position within the apparatus and a risky event ethogram capturing selected behaviours associated with physical challenge. By comparing the behaviour of animals during the natural peak of play behaviour with animals beyond this developmental period, we aimed to evaluate whether engagement with elevated and unstable environments reflects play-related processes rather than environmental complexity alone. This study provides a behavioural framework for investigating risky play-like behaviour in rodents and establishes a foundation for future studies examining its developmental and neurobiological significance.

## Introduction

Play is observed in multiple species across the animal kingdom, and is particularly conserved in mammals (Burghardt, 2005; Fagen, 1981). Across species, play emerges early in development, occurs in the absence of immediate survival benefits and often involves behaviours that appear energetically costly, inefficient or even potentially dangerous. Despite these apparent costs, play has been retained throughout evolution, suggesting that it contributes important developmental functions that outweigh its immediate disadvantages (Pellis & Pellis, 2009; Spinka et al., 2001).

Animal play is commonly divided into three broad categories: locomotor play, object play and social play (Burghardt, 2005; Pellegrini & Smith, 1998). Locomotor play includes vigorous running, jumping, climbing and other exaggerated movements, object play involves interactions with inanimate features of the environment and social play encompasses playful interactions between individuals. Although these forms of play differ behaviourally, they share several defining characteristics, including intrinsic motivation, behavioural flexibility, repetition without stereotypy and modification of otherwise functional behaviours (Burghardt, 2005).

Experimental studies have demonstrated that juvenile play, peaking between postnatal day (PND) 21 and 42 in rats, contributes to the development of social competence, emotional regulation, behavioural flexibility and executive functioning (for review see Achterberg & Vanderschuren, 2023; Bell et al., 2010; Pellis & Pellis, 2009). Several studies have revealed anomalies in anxiety, stress and social behaviour in rats that were deprived of social play behaviour (Baarendse et al., 2013; Bell et al., 2010; Bijlsma et al., 2022; Hol et al., 1999; Lukkes et al., 2009; van den Berg et al., 1999; Von Frijtag et al., 2002). These findings suggest that play actively contributes to healthy development.

Although considerable progress has been made in identifying developmental consequences of play, the mechanisms through which play exerts these effects remain incompletely understood. A recurring theme across several theoretical accounts is that play exposes animals to situations characterized by uncertainty, instability, rapid behavioural adjustments and temporary loss of control (Pellis & Pellis, 2009; Spinka et al., 2001). According to the *training-for-the-unexpected* hypothesis, play allows animals to experience unexpected events in a relatively safe context, thereby promoting the development of flexible behavioural responses to future challenges (Spinka et al., 2001). On the other hand, play offers ample opportunities to encounter and manage uncertainty. Indeed, mammals voluntarily engage in vigorous locomotion, unstable postures, rapid directional changes, physical collisions and situations in which balance or behavioural control may be temporarily compromised (see e.g. Byers & Walker, 1995; Pellis & Pellis, 2009). These behaviours often involve a possibility of mistakes, falls or minor injury, yet animals repeatedly seek out such experiences even when less challenging alternatives are available. Such observations raise the possibility that voluntary exposure to manageable forms of physical challenge may itself constitute an important feature of play. Indeed, the first studies to capture risky play opportunities emphasize the long-term impact of such risky play encounters on the development of both brain and behaviour (e.g. Bijlsma et al., 2024).

Related to this idea, Sandseter (2007) introduced the concept of risky play in developmental research in children to describe thrilling and exciting forms of play involving uncertainty and a possibility of physical injury. Risky play includes activities involving height, speed, unstable surfaces, rough-and-tumble interactions and situations in which children experience temporary loss of control. Importantly, risky play is not defined by actual harm but by voluntary engagement with manageable risk. Voluntary engagement with physical challenge represents a previously overlooked developmental factor. However, increasing evidence suggests that opportunities for risky play contribute to motor development, risk assessment, resilience, emotional regulation and adaptive coping with uncertainty (Brussoni et al., 2015; Brussoni et al., 2012; Sandseter & Kennair, 2011).

The concept of risky play is particularly interesting from a comparative perspective because many of its defining features closely resemble characteristics already emphasized in theories of animal play. Climbing elevated structures, traversing unstable substrates, jumping across gaps, engaging in rough-and-tumble interactions and experiencing temporary loss of balance are all behaviours frequently observed in juvenile mammals. However, translating the concept of risky play from children to non-human animals presents substantial conceptual challenges. Behaviours such as climbing, balancing and jumping are naturally occurring components of the rat behavioural repertoire and therefore cannot automatically be classified as risky play. Furthermore, subjective experiences such as thrill, excitement, or perceived danger cannot be directly measured in animals. Consequently, any operationalization of risky play in rodents must be based on observable behavioural characteristics rather than inferred psychological states.

The present study aimed to establish an ethological framework for identification and quantification risky play-like behaviour in rats. Social play is most pronounced during the juvenile period (Panksepp, 1981; Vanderschuren et al., 2016), but little is known about the ontology of other forms of play. Therefore, both juvenile and young adult animals were repeatedly exposed to a custom-designed high enclosure containing elevated elements, which were either stable or unstable, providing opportunities for climbing, balancing, hanging, jumping and social interaction. We quantified location use, risky behavioural events and location-specific disbalance rates to determine whether animals voluntarily engaged with physically challenging features of the environment. We predicted that behaviours associated with risky play would be most prominent during the juvenile period of peak play expression. By providing a behavioural framework for identifying risky play-like behaviour in rodents, this study establishes a foundation for future investigations into the developmental and neurobiological significance of risky play in animal models.

## Methods

### Animals

Thirty-six male Lister Hooded rats (*Rattus norvegicus*) were obtained from Charles River Laboratories (Sulzfeld, Germany) and housed at the animal facility of Utrecht University. Upon arrival at postnatal day (PND) 12, animals were housed in litters together with their nursing dam. Following weaning (PND23), one animal was excluded due to being a female. The remaining rats were pair-housed in standard Type IV Makrolon cages under a reversed 12:12 h light–dark cycle (lights off at 07:00 h) with ad libitum access to food and water. Two developmental cohorts were studied. Juvenile rats (n = 23) were exposed to the risky play environment between approximately PND21 and PND42, corresponding to the developmental period during which play behaviour reaches peak expression in rats. Young adult rats (*n* = 12) were exposed between approximately PND50 and PND71, after the juvenile peak in play behaviour. All Experimental procedures were approved by the Animal Ethics Committee of Utrecht University and the Dutch Central Authority for Scientific Procedures on Animals (CCD), and conducted in agreement with Dutch (Wet op de Dierproeven 1996; Herziene Wet op de Dierproeven 2014) and European legislation (Guideline 86/609/EEC; Directive 2010/63/EU).

### Experimental Design

Animals were exposed to a custom-designed risky play environment for 30 min per weekday throughout the experimental period (Figure 1). Rats were always placed into the apparatus together with their cage mate to preserve the social context in which play behaviour naturally occurs. To characterize both interactions with the environment and behavioural changes following repeated exposure, video recordings from the first and final exposure sessions were selected for behavioural analysis.

**Figure 1.**
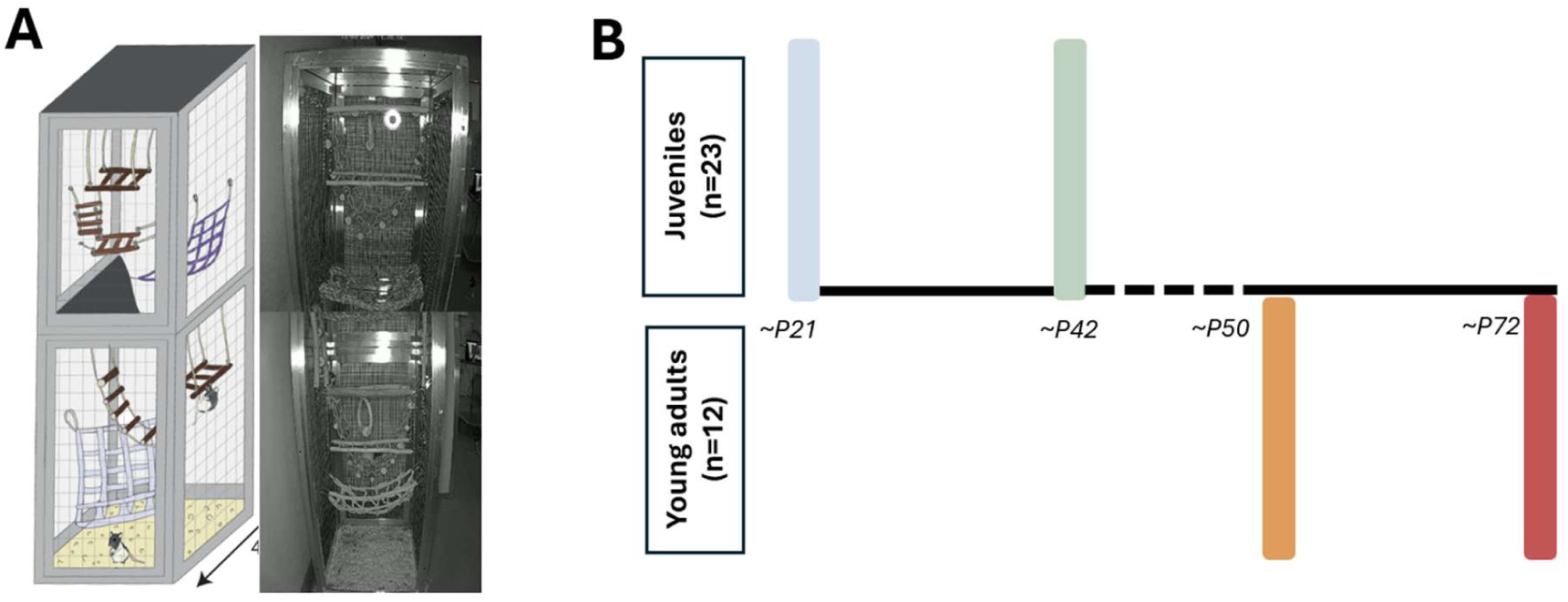
Experimental design. **A** shows the schematic diagram of the risky play setup with a bottom and upper part where both stable and wobbly elements are placed. **B** shows the time-window for the risky play sessions for juvenile rats (∼PND 21-42) and young adult rats (∼PND50-72), respectively. The behavioural observations were made at the first and last day of this time-window; the colours are aligned with the figures in the results section.

### Risky Play Environment

The risky play environment consisted of a vertically structured enclosure (40 × 40 × 150 cm) divided into an upper and lower level. The apparatus was designed to provide opportunities for voluntary engagement with elevated locations and unstable elements. Both levels contained identical structural elements, including a perforated mat, a bridge and a horizontal ladder. Elements were classified as either *steady* or *wobbly*. Steady elements remained stationary, whereas wobbly elements moved in response to the animal’s body weight and movement. The inclusion of identical structures on both the lower and upper level allowed for independent examination of the effects of elevation and instability (Figure 1A). The risky play environment further provided opportunities for climbing, balancing, hanging, jumping and social interaction. All sessions were conducted during the dark phase between 09:00 and 12:00 under dim red illumination and were video-recorded using BASCOM cameras.

### Behavioural Observations

Video recordings were analysed using Observer XT 17 (Noldus Information Technology, Wageningen, The Netherlands). Two ethograms were developed for the present study: a location ethogram and a risky event ethogram. The location ethogram (see Table 2) was used to score the animal’s position within the apparatus, whereas the risky event ethogram (see Table 1) was used to score selected behaviours associated with risky play-like behaviour.

**Table 1.** Behavioural definitions used in the risky event ethogram.

| Event type | Behaviour | Definition |
| --- | --- | --- |
| Point | Jump | Any event in which an animal voluntarily propelled itself from one location to another while all four paws simultaneously lost contact with the supporting surface. |
| Point | Disbalance | Any observable disruption of postural stability resulting in a sudden loss of balance. Disbalance events were further classified as either *Correction* (balance successfully restored) or *Fall* (instability resulting in uncontrolled displacement or descent). |
| State | Hanging | Any event in which part of the animal's body was unsupported and suspended in space while the animal remained attached to a structure by one or more limbs. Hanging could occur with either the forequarters or hindquarters unsupported, provided that the unsupported body region was clearly elevated from surrounding surfaces and maintained elevated against gravity. Animals suspended beneath a platform or mesh structure were also scored as hanging. |
| State | Social Play | A play bout consisting of pouncing and pinning interactions between cage mates. A bout could contain a single pounce or pin, or multiple consecutive pouncing and pinning interactions. Consecutive interactions were considered part of the same bout until the animals disengaged and no longer interacted through pouncing or pinning behaviour. |

**Table 2.** Conceptual framework underlying location weights.

| **Location Category** | **Height** | **Instability** | **Assigned Weight** | **Rationale** |
| --- | --- | --- | --- | --- |
| Bottom–Steady | No | No | 1 | Baseline condition with neither elevation nor instability. |
| Bottom–Wobbly | No | Yes | 2 | Contains instability but no elevation. |
| Upper–Steady | Yes | No | 2 | Contains elevation but no instability. |
| Upper–Wobbly | Yes | Yes | 4 | Combines elevation and instability, representing the highest level of physical challenge within the apparatus. |

*Location weights were assigned based on the assumption that elevation and instability constitute independent dimensions of physical challenge. Locations containing one challenge dimension received an intermediate score, whereas locations combining both dimensions received the highest score.*

### Observer Training and Reliability

Because no established ethogram exists for quantifying risky play-like behaviour in rats, considerable effort was devoted to observer training and reliability assessment. The initial development of the ethograms involved independent scoring of the same recordings from another study by at least two observers. Following scoring, all identified events were systematically compared and reviewed. Discrepancies were discussed on an event-by-event basis until consensus was reached regarding behavioural definitions, inclusion criteria and timing of scoring. This process resulted in a dataset that served as a reference for subsequent observer training. New observers were trained using this reference material and were required to achieve a minimum inter-rater reliability score of 80% before scoring experimental recordings independently. Training continued until this criterion was met consistently across at least three training recordings. Behavioural scoring was conducted over an extended period, therefore periodic reliability checks were incorporated throughout the scoring phase.

### Location Ethogram

Locations were classified according to both vertical position and element stability, resulting in four mutually exclusive states: Bottom–Steady, Bottom–Wobbly, Upper–Steady, Upper–Wobbly. Location was determined by the position of the hind paws, as these represent the primary point of support during locomotion. A transition between locations was scored when both hind paws entered a new location category. During hanging events, if the hind paws were no longer in contact with a supporting surface, location was assigned based on the structure to which the animal remained attached.

### Risky Event Ethogram

Four behavioural categories were scored: Jump, Hanging, Social Play and Disbalance (Table 1). Jump and Disbalance were scored as point events, whereas Hanging and Social Play were scored as state events to also capture their duration.

### Risky Play Score

No established quantitative framework currently exists for measuring risky play in rodents, therefore a cumulative Risky Play (RP) score was developed to integrate information from both the location and risky event ethograms. Ordinal weights were assigned to location categories and behavioural events based on their presumed level of physical challenge within the context of the apparatus. The weighting scheme was designed to provide a first approximation of engagement with elevated, unstable and physically challenging aspects of the environment. For location categories, Bottom–Steady, Bottom–Wobbly, Upper–Steady and Upper–Wobbly were assigned values of 1, 2, 2 and 4, respectively (Table 2). The weighting scheme reflected the assumption that elevation and instability represent two independent dimensions of physical challenge. Locations containing either elevation or instability received an intermediate score, whereas locations combining both characteristics received the highest score. For behavioural events, Jump, Social Play, Hanging and Disbalance were assigned values of 1, 2, 3 and 4, respectively (Table 3). This ranking was based on the presumed degree of physical challenge and postural instability associated with each behaviour. Jumping was considered the least challenging event because it reflects voluntary locomotor activity without an observable loss of control. Social play was assigned a higher value because interactions with a play partner introduce behavioural unpredictability. Hanging was ranked higher because part of the body is unsupported and suspended against gravity. Disbalance received the highest value because it represents an observable disruption of postural stability and therefore the behaviour most directly associated with temporary loss of control. The weighting scheme should not be interpreted as an objective measure of risk. Rather, it represents an a priori ranking intended to facilitate exploratory quantification of risky play-like behaviour. Future work may refine or validate this framework using empirical measures of motor challenge, energetic cost or behavioural consequences.

**Table 3.** Conceptual framework underlying behavioural weights.

| Behaviour | Unpredictability | Postural Challenge | Loss of Control | Assigned Weight | Rationale |
| --- | --- | --- | --- | --- | --- |
| Jump | Low | Moderate | No | 1 | Voluntary locomotor challenge without observable instability or loss of control. |
| Social Play | Moderate | Moderate | Limited | 2 | Introduces behavioural unpredictability through interaction with a play partner. |
| Hanging | Low | High | Potential | 3 | Requires active support of the body while part of the body remains unsupported against gravity. |
| Disbalance | High | High | Yes | 4 | Represents an observable disruption of postural stability and the behaviour most directly associated with temporary loss of control. |

*Behavioural weights were assigned according to the presumed degree of physical challenge and instability associated with each behaviour. These values were intended as an exploratory ranking rather than a validated measure of risk.*

### Data Extraction and Processing

Behavioral data were exported from The Observer XT 17 and processed with custom MATLAB R2025b (MathWorks) scripts. To equalize observation time for all animals, we used a strict 22-minute (*1320 seconds*) cut off time. Next, we combined data from two different ethograms: the continuous location ethogram (table 2) and a discrete risky event ethogram (table 1). For location data, we calculated the percentage of time spent in four different cage zones based on height and stability (*Bottom-Steady*, *Bottom-Wobbly*, *Upper-Steady*, *Upper-Wobbly*). For the risky event ethogram, we calculated the frequencies and durations of specific behaviors (*Jumping*, *Disbalance*, *Hanging*, and *Social Play*). Disbalance events were also split into *Corrections* (recovering balance) and *Falls* (losing balance). To find out the exact location where a risky behavior happened, a custom-written MATLAB script aligned the timestamps of both datasets. This script automatically added the cage location to each behavioral event. After this, we calculated three different scores: 1) Location Score: Calculated per zone by multiplying the time spent in that zone by a risk weight (*Bottom-Steady = 1; Bottom-Wobbly = 2; Upper-Steady = 2; Upper-Wobbly = 4*). 2) Behavior Score: Assigned based on the risk of the behavior itself (*Jumping = 1; Social Play = 2; Hanging = 3; Disbalance modifiers = 4*). 3) Total risk score: A combined score for both location and behavior. For point events (*Jumping*, *Corrections*, *Falls*), the score is the Location Score multiplied by the Behavior Score. For state events (*Hanging*, *Social Play*), the score is the Location Score multiplied by the Behavior Score and the exact duration of the event. To assess whether different locations varied in their associated challenge, disbalance events were normalized to the amount of time spent within each location category. This yielded a location-specific disbalance rate expressed as the number of disbalance events per unit time. Locations associated with higher failure rates were interpreted as imposing greater motor challenge.

To avoid missing values when rats did not show a behavior or location, the frequency and duration were explicitly filled with a zero instead of leaving them blank.

All parameters were extracted separately for the first session (*PND21 for juveniles, PND50 for young adults*) and the final session (*PND42 for juveniles, PND70 for young adults*). Transition matrices were made to show the movements between the four zones visually, but these were not used for statistics.

### Statistical Analysis

Statistical analyses were done in RStudio (version 2025.05.0) with significance set at *p* < 0.05. We used mixed-effects models, with Rat ID as a random intercept in all models to account for the repeated measurements. Main effects and interactions were tested with Type III Wald chi-square tests. *Post-hoc* comparisons were performed using estimated marginal means with Bonferroni corrections.

Continuous variables, like total time in location zones, Location and Risk scores, latency after the first disbalance, and mean inter-event intervals, were analyzed with Linear Mixed Models (LMMs). To visualize changes over time during a session, location and risk scores were plotted per minute. However, for the statistical analysis, data were grouped into 5-minute time bins and compared with each other, resulting in 4 complete time bins per session. For spatial allocation, Age, Session, and Location were the fixed factors. For the 5-minute bins, we ran separate LMMs for juveniles and young adults with Session, TimeBin, and their interaction as fixed factors. For the kinetic parameters (latency and inter-event intervals between risky events), sessions with zero events were treated as missing data and left out of those specific models.

Discrete count data, such as frequencies of zone entries, total risky events, individual risky behaviors, and disbalance modifiers (correction or fall), were analyzed with Generalized Linear Mixed Models (GLMMs) using a Poisson distribution and a log-link function. Zone transition matrices were only used for visual description and were not tested statistically.

## Results

### Spatial Dynamics and Engagement with Height and Instability Across Development

To evaluate how animals interacted with the play environment, we analysed their spatial allocation, location scores, and visualized the location transitions (Figure 2). For the total time spent in different locations, a significant three-way interaction between Age, Session, and Location was observed (*χ*^2^ (3) = 23.12, *p* < 0.0001; Figure 2A). Post-hoc tests showed that both juvenile and young adult rats spent significantly less time in the safest zone (Bottom-Steady) during the final session compared to their first exposure (Juveniles: *p* < 0.0001; Young Adults: *p* < 0.0001). At the same time, the time spent in the more challenging locations (B-W, U-S, and U-W) did not show significant changes between sessions (all *p* > 0.15). This indicates that the general preference for specific locations did not change much over time for either age group. This pattern was further supported by entry frequencies, which showed a significant three-way interaction (*χ*^2^ (3) = 8.10, *p* = 0.044; Figure 2B). Young adults increased their entries into the baseline Bottom-Steady zone during the final session (*p* < 0.0001), while juveniles maintained an active and evenly distributed entry frequency across all structural zones without significant session shifts (*p* = 0.057). For the Location score, the interaction between Age and Session was not significant (*χ*^2^ (1) = 0.04, *p* = 0.850; Figure 2C). Instead, a strong main effect of Session was found (*χ*^2^ (1) = 24.42, *p* < 0.0001), demonstrating that both age groups significantly increased their total cumulative Location Scores by the end of the study (Juveniles: *p* < 0.0001; Young Adults: *p* = 0.0040).

**Figure 2.**
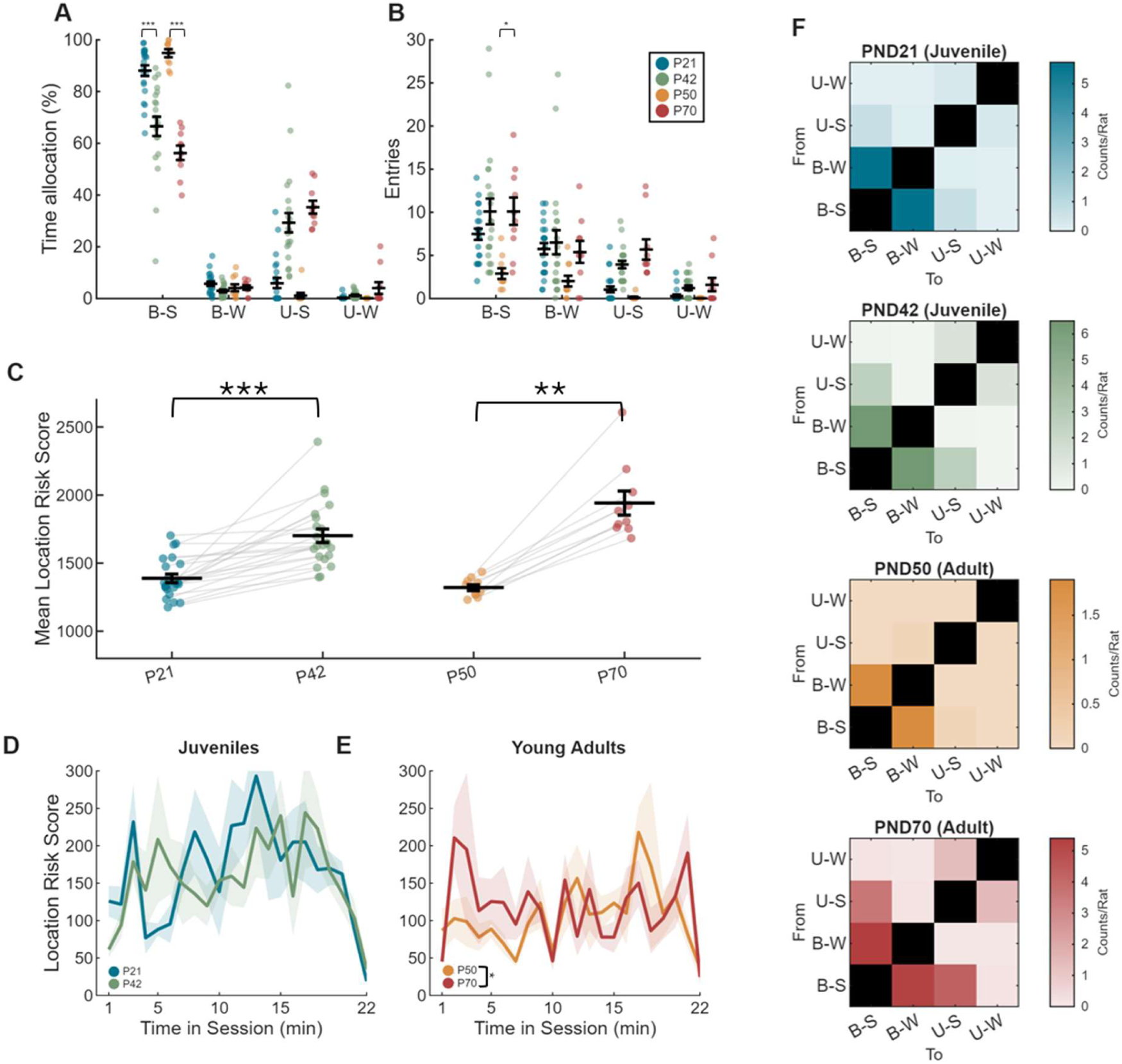
Spatial dynamics, location scores, and movement transitions during the first and final sessions in the playground environment. (A) Percentage of total duration spent in the four location zones: Bottom-Steady (B-S), Bottom-Wobbly (B-W), Upper-Steady (U-S), and Upper-Wobbly (U-W). Dots represent individual scores; error bars show mean ± SEM. Data points are organized by developmental stage (Juveniles and Young Adults) and testing session. (B) Frequency, shown as the average number of entries into each location zone per session. (C) Cumulative location scores across developmental cohorts, calculated as the sum of the pre-assigned risk weight of each location multiplied by the exact duration spent in that zone. Light grey lines track within-subject shifts between sessions; horizontal bars show group means ± SEM. Within-session exploration kinetics illustrated by cumulative location Scores per minute for (D) juvenile and (E) young adult rats. Solid lines show mean location scores and shaded areas represent SEM. (F) Spatial transition matrices visually showing the average frequency of sequential movements between the four location zones (From/To) per cohort and per session. Darker colors indicate a higher frequency of transitions between specific zones, as defined by the respective color scales.

**Figure 3.**
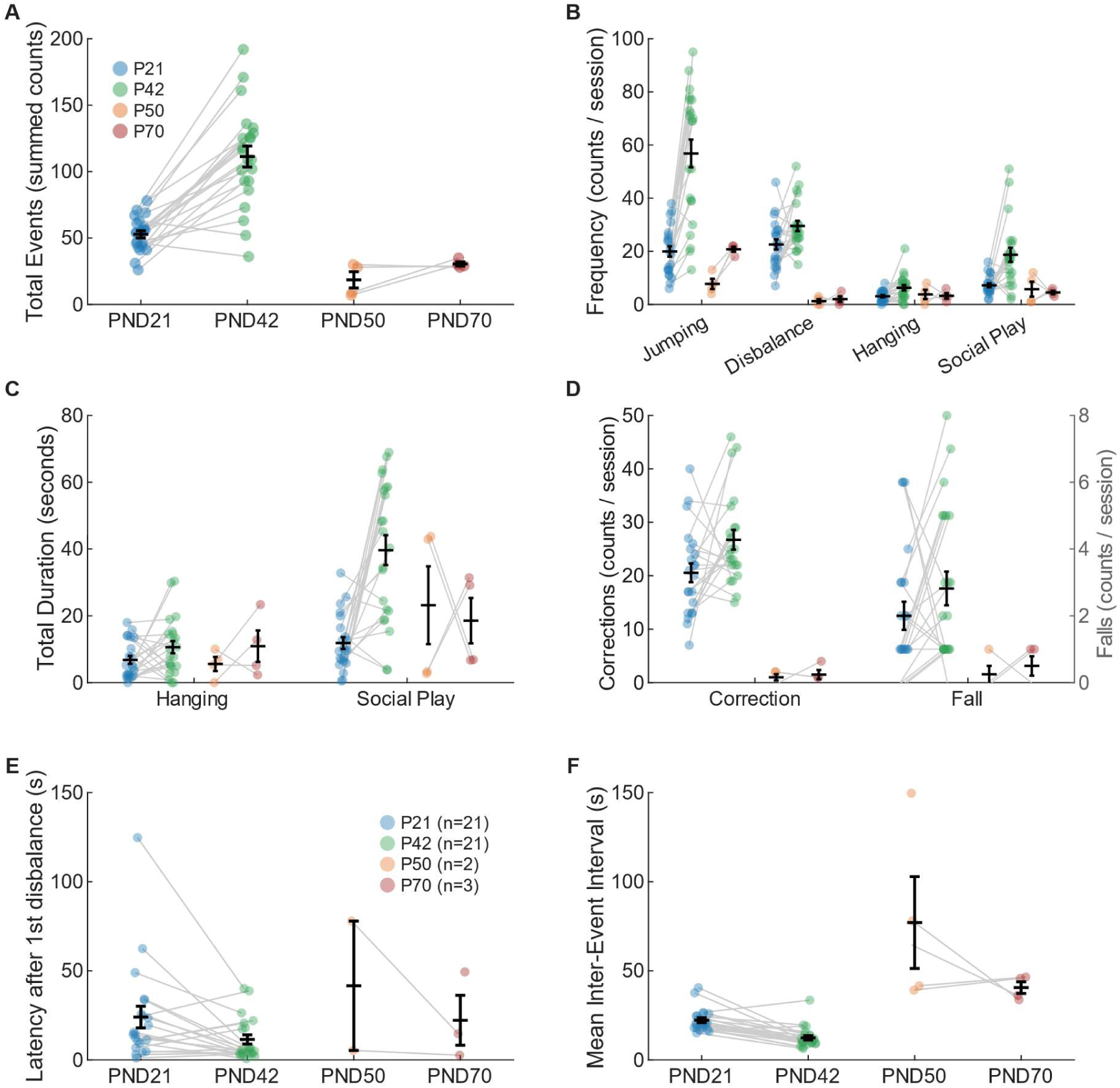
Shifts in the structure, composition, and temporal kinetics of selected behaviors in the setup. Detailed behavioral profiling of risk-taking patterns during play sessions across juvenile (PND21, PND42) and adult (PND50, PND70) rats. (A) Total combined frequency of all recorded risky events per session, with grey lines representing paired individual trajectories across sessions. (B) Frequencies of discrete motor patterns including Jumping, Disbalance, Hanging, and Social Play. (C) Cumulative session duration for state behaviors (Hanging and Social Play, total duration, not average anymore). (D) Frequency of disbalance modifiers, displaying disbalance → corrections (left y-axis) and disbalance → falls (right y-axis). (E) Latency between the first disbalance and the next subsequent behavioral event. Sample sizes per session are updated in the legend to reflect only those animals that experienced a disbalance followed by further behavioral activity within the session (F) Mean inter-event interval representing the average time window between consecutive risky behaviors (not specific, any event). Data points represent individual subjects (colored circles) plotted with group means ± SEM (black crossbars). Young adult data not yet completed.

Clear differences between the age groups appeared when looking at the within-session kinetics. For juvenile rats, the pattern of location score accumulation was consistent across time, as shown by a non-significant Session by TimeBin interaction (*χ*^2^ (5) = 3.00, *p* = 0.700; Figure 2D). Juveniles showed a strong main effect of TimeBin (*χ*^2^ (5) = 193.37, *p* < 0.0001), which was driven by a peak during the first half of the trials. A session-driven increase occurred only during the first 5 minutes (TimeBin 1: *p* = 0.030). In contrast, young adult rats showed a dynamic shift, resulting in a significant Session by TimeBin interaction (*χ*^2^ (5) = 13.17, *p* = 0.0218; Figure 2E). While young adults started with very low activity in the first session, they increased their Location Scores during the final session across almost all early and middle time blocks. This activity peaked during the first 5 minutes (TimeBin 1: *p* < 0.0001) and remained high from minutes 5 to 20 (TimeBins 2–4: *p* < 0.02) before decreasing back to baseline at the end of the session (TimeBin 6: *p* = 1.000). Finally, the spatial transition matrices (Figure 2F) visually show these different strategic behaviors. Juveniles displayed a highly distributed, complex transition network from their first exposure, moving between all four structural levels and navigating directly toward the upper wobbly elements. On the other hand, young adults displayed a restricted movement strategy during their first session, moving almost exclusively between the two bottom locations, and the young adults expanded their transitions toward the upper parts of the setup only after full cross-session habituation.

### Quantification of behavioral profiles

*The dataset for the young adults is not yet complete, but it has already been visualised. The scripts for the statistical analysis are also ready; the dataset just needs to be completed. I now have n=4 for both sessions of the young adults (total: n=12, so there are still 8 rats × 2 sessions to score for the risky events ethogram). The statistics below relate only to the juveniles (between-sessions), but ultimately I also want to make a comparison with the young adults. Figures have already been plotted, even though n is low for the young adults (with regard to risky events)*.

For the total number of behavioral events, the main model revealed a significant main effect of Session (*χ*^2^ (1) = 11.56, *p* = 0.0007). Post-hoc comparisons focused on the juvenile cohort demonstrated a significant increase in the total number of events from the first to the final session (*p* < 0.001). When analyzing the frequencies of specific behavioral categories, the occurrence of *Jumping* was significantly affected by Session (*χ*^2^ (1) = 22.07, *p* < 0.001), with juveniles showing a significant increase in jumping frequency between the first and final session (*p* < 0.001). For *Disbalance* frequency, there was no overall main effect of Session (*χ*^2^ (1) = 0.68, *p* = 0.409), but pairwise comparisons within the juvenile group revealed a significant increase in disbalance events from the first to the final session (*p* < 0.001). The frequency of *Hanging* showed a significant Age x Session interaction (*χ*^2^ (1) = 4.45, *p* = 0.035), and post-hoc tests confirmed that juveniles performed hanging behavior significantly more often during the final session compared to the first session (*p* < 0.001). Similarly, *Social Play* frequency displayed a significant Age x Session interaction (*χ*^2^ (1) = 13.56, *p* = 0.0002), driven by a significant increase in play behavior frequency in juveniles from the first to the final session (*p* < 0.001). The time the animals spent *Hanging* did not change significantly over sessions (*χ*^2^= 1.07, *p* = 0.301). In contrast, the total duration of *Social Play* showed a significant Age x Session interaction (*χ*^2^ (1) = 8.58, *p* = 0.003), with juveniles spending significantly more time engaged in social play during the final session than during the first session (*p* < 0.001). Regarding disbalance modifiers, the frequency of *Corrections* did not show a main effect of Session (*χ*^2^ (1) = 0.40, *p* = 0.529), yet pairwise comparisons indicated that juveniles experienced significantly more corrections in the final session (*p* < 0.001), while the frequency of *Falls* did not change significantly over time (*χ*^2^ (1) = 0.32, *p* = 0.571) or within the juvenile cohort (*p* = 0.082). Finally, looking at behavioral kinetics, the latency to the next event after the first disbalance showed no global session effect (*χ*^2^ (1) = 1.38, *p* = 0.239), but within-group comparisons revealed a significant decrease in latency for juveniles between the first and final session (*p* = 0.045). The mean inter-event interval was significantly affected by Session (*χ*^2^ (1) = 13.25, *p* = 0.0003) and showed a significant Age x Session interaction (*χ*^2^ (1) = 5.97, *p* = 0.015), with juveniles displaying a significant reduction in the average time between consecutive events from the first to the final session (*p* = 0.029).

### Development of risky play-like behaviors and spatial distribution in the cage

To understand how risky play behavior and its spatial dynamics change as animals grow up, we mapped the frequency of specific risky behaviors across different cage locations for each age group (Figure 4A– D). In young juvenile rats (PND21), risky play mostly consisted of jumping and social play, which took place almost completely in the steady bottom part of the cage (Figure 4A). At PND42, these older juveniles still preferred the steady bottom zone, but they also started to expand their behavior towards other zones in the cages. They showed an increase in behaviors that are associated with balance, like hanging and making disbalance-corrections, in both the steady and the wobbly upper parts of the cage (Figure 4B). We also tracked a separate group of rats longitudinally during young adulthood to determine whether these behaviors are displayed and evolve in a comparable or different fashion at ∼PND50 and ∼PND72 (Figure 4C–D). Interestingly, at the first session (PND50), risky play was almost completely absent (Figure 4C; This may still change – the dataset is not complete!). However, by PND70, risky play behavior returned and showed a change in location. Jumping on the steady bottom was still the most common behavior, but adult rats at this stage expanded their movement to higher areas. They showed a increase in jumping and social play in the wobbly bottom and steady upper zones (Figure 4D; subject to change).

**Figure 4.**
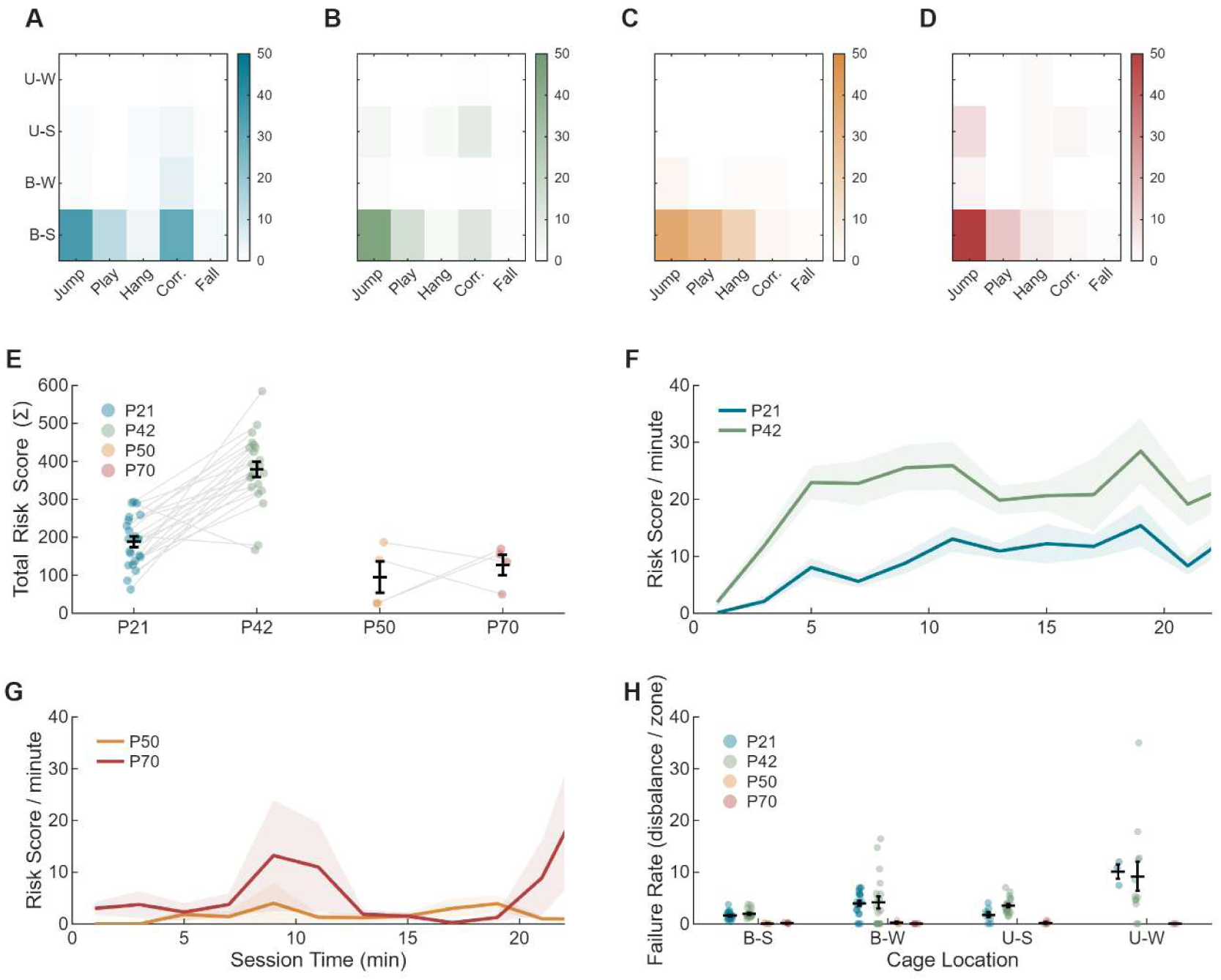
Behavioral repertoires, temporal kinetics, and spatial distribution of risky events/score across development. (A– D) Heatmaps showing the total frequency of specific risky events across different cage locations (B-S: bottom-steady; B-W: bottom-wobbly; U-S: upper-steady; U-W: upper-wobbly) for (A) PND21 juveniles, (B) PND42 juveniles, (C) PND50 adults, and (D) PND70 adults. Color intensity represents the total event count. (E) Total risk score per individual rat across age groups; grey lines connect longitudinal sessions within the juvenile (P21 to P42) and adult (P50 to P70) cohorts. (F–G) Temporal kinetics showing the mean (± SEM) risk score per minute over the session for (F) juveniles and (G) adults. (H) Failure rate (number of disbalance events divided by location duration) across the four cage locations. For panels E and H, individual dots represent single animals, and black error bars represent the mean ± SEM. Data young adults niet complete!

To statistically test whether risk behaviors changed over time, we analyzed the total risk score across behaviors (Figure 4E). The linear mixed-effects model showed a significant interaction between Age and Session (*χ*^2^ (1) = 6.53, *p* = 0.011). This was driven by a change during development in the juvenile group. Specifically, juvenile rats showed a strong and highly significant increase in their total risk score from their first session on PND21 to their final session on PND42 (post hoc contrast: *t*(26.6) = -7.85, *p* < 0.0001; Figure 4E). Add YA statistics when the dataset is complete. Next, we looked at the temporal kinetics (the risk score per minute) during the sessions (Figure 4F–G). This minute-by-minute rate of the risk was significantly influenced by an interaction between Session and the time bins, which depended on the Age Cass (Age × Session × TimeBin interaction: *χ*^2^ (1) = 0.09, *p* = 0.765; Age Class × TimeBin interaction: *χ*^2^ (1) = 4.50, *p* = 0.034). Within the juvenile group, the pattern over time changed significantly between sessions (post hoc slope contrast: *t*(1114) = 2.41, *p* = 0.016). On PND21, juveniles had a flat and stable rate of play throughout the session (slope trend = 0.01). On PND42, they showed a more dynamic pattern with an early peak followed by a slow decrease toward the end of the session (slope trend = -0.33; Figure 4F). Finally, we looked at how challenging the different locations of the cage were by analyzing the rate of behavioral failures (Failure Rate; Figure 4H). The linear mixed model showed a very strong main effect of Location (*χ*^2^ (3) = 18.75, *p* = 0.0003), while there was no significant effect of Session (*χ*^2^ (1) = 0.11, *p* = 0.737) or interaction between Session and Location (*χ*^2^ (3) = 1.09, *p* = 0.780). Post hoc tests showed that the wobbly elements caused significantly more behavioral failures than the steady elements. Specifically, the failure rate on the wobbly bottom (B-W) was significantly higher than on the steady bottom (B-S; *t*(102) = -2.96, *p* = 0.023). This effect was even stronger for the upper wobbly platform (U-W), which showed an increase in failures compared to both the steady bottom (*t*(112) = -6.28, *p* < 0.0001) and the steady upper platform (U-S; *t*(110) = -5.19, *p* < 0.0001; Figure 4H). These results show that the wobbly locations are highly challenging on their own. This difficulty may explain why these specific zones are less visited in the behavioral matrices (Figure 4A–B); because these elements are highly unstable, even a very short visit or a small number of actions immediately/quickly results in a strong challenge to the animal’s balance, meaning they do not need to spend a lot of time there to experience risk. Importantly, the lack of a Session effect confirms that when older juveniles play in more challenging ways on PND42, it does not lead to more motor mistakes compared to PND21. // Describe the young adults data once the dataset is complete.

## Discussion

This study introduces a novel framework to identify and quantify risky play-like behavior in laboratory rats, demonstrating that voluntary engagement with physical challenge is a regulated process rather than a passive response to environmental complexity. By tracking two separate developmental cohorts longitudinally, observing the juvenile cohort at the onset (PND21) and the end (PND42) of their natural play peak, and a separate young adult cohort at two post-play peak stages (PND50 and PND70; first and last play session), we mapped how rats manage uncertainty over time. We found a difference in how these two cohorts interacted with the play setup. While the younger juveniles initially confined their behavioral activities almost exclusively to the stable level (location: bottom-steady), tracking this same cohort to PND42 revealed a significant expansion of their spatial and behavioral repertoire into elevated and unstable zones in the setup. Crucially, our assessment of location-specific disbalance rates revealed that the wobbly platforms imposed a higher (motor) challenge level, significantly increasing disbalance events compared to the stable elements. Despite the high risk associated with the wobbly platforms, older juveniles chose to interact with these features without experiencing more falls. This may suggest that being exposed to risky playful elements from an early age calibrates balance and motor skills during the peak play period, showing that physical risk-taking may have a function in their motoric development.

The behavioral expansion in the risky play cages observed between PND21 and PND42 highlights a critical developmental window during which juvenile mammals play actively. This phenomenon aligns closely with the *training-for-the-unexpected* hypothesis formulated by Spinka et al. (2001), which posits that the primary function of play is to expose the animal to unexpected loss of control to practice emotional and motor recovery mechanisms. By deliberately executing balance-challenging movements like hanging and making sudden corrections on highly unstable elements, PND42 juvenile rats maximize sensory-motor feedback during a phase of intense neural development. This active engagement with environmental unpredictability mirrors the concepts of motor automation and physical auditing detailed by Byers and Walker (1995), who argued that locomotor play is timed to coincide with the structural development of cerebellum circuitry. Our data closely supports this idea, although more interactions with the unstable elements led to a higher rate of disbalance events, it did not result in more actual falls. This demonstrates that even though the wobbly platforms were physically challenging, the older juveniles were capable of quickly correcting their movements and recovering their balance. They appear to use their playful motivational state to test the boundaries of their physical environment and their own motor limits, successfully balancing the drive for novelty with the management of immediate physical harm.

Crucially, the comparison with our young adult cohort provides a developmental control that reinforces the status of these juvenile activities as play, considering that young adult animals are less motivated to engage in play. Although both cohorts were exposed to the exact same environment, the young adults tracked longitudinally at PND50 and PND70 exhibited lower total risk scores and displayed less challenging behavioral events compared to the juvenile cohort. Adult rats are larger, stronger, and physically more competent than juveniles, therefore their avoidance or highly restricted interaction with the wobbly and elevated zones demonstrates that environmental complexity alone does not trigger these challenging behaviors. In this context, the young adult cohort may be considered as a negative control for non-playful exploration: they occupy the same space but apparently lack the age-specific developmental drive to test their physical boundaries. When young adult animals were exposed to the risky play setup for the first time, they did interact with the setup but their interactions were mostly limited to basic exploration of the lower levels (PND50). They did not show the high rates of jumping, hanging, or social play that we observed in the juvenile cohort (The dataset is not yet complete…). Besides playing, the few social interactions we saw in young adults were dominance behaviors on the cage floor, a type of behavior not included in the ethogram (mostly boxing, not followed by pounces of pins) that was not observed in the juvenile animals. The developmental decline in physical risk-seeking fits perfectly within the wider trajectory of rodent behavior documented by Panksepp (1981) and reviewed by Vanderschuren et al. (2016), where older animals (post juvenile period) shift their energy budgets away from highly flexible, peer-directed play toward predictable, energy-conserving, and utilitarian activities such as foraging, territorial maintenance, and established social hierarchies.

While this study establishes a framework for measuring risky play-like behavior, several methodological limitations regarding our ethogram designs must be addressed. Within the location ethogram, the *Bottom-Steady* category currently encompasses both the floor of the cage and the low-level stationary platforms. In retrospect, scoring the cage floor as a completely separate zone would have provided a higher resolution regarding the animals’ choices with regards to location in the set-up. Differentiating the floor from slightly elevated but stable elements would allow us to verify whether animals were truly seeking vertical height or simply remaining on the lowest horizontal plane. In addition to refining the ethograms, future studies should expand the developmental scope by testing a third, fully mature adult cohort beyond PND90. In the current design, our independent young adult cohort was first exposed to the apparatus at PND50. While PND50 in rats is just beyond the classic juvenile play peak (which rapidly declines after PND42), PND50 rats are biologically still in a transitional phase of late adolescence or early adulthood (Spear, 2000), which may translate into a lower yet existent exploratory drive. Therefore, future experiments should consider to include a separate cohort of fully mature adults (>PND90) as fully adult animals will have completely stabilized their energy budgets, fully consolidated their prefrontal cortical circuitry, and established social hierarchies (Bell et al., 2010; Newman, 1999). Adult rats would likely utilize the apparatus in a different manner than juveniles, thereby aiding in isolating play-related processes from adult behaviors even further.

The findings also have broader implications, for example for the welfare, and cognitive development of children and (laboratory) animals,. Standard laboratory caging is generally restrictive, offering animals virtually no opportunities for vertical exploration, climbing, or interaction with unstable surfaces. Some laboratories implemented multi-level cages, adhering better to the natural needs of rodents. By showing that juvenile rats actively seek out and manage physical uncertainty when given the opportunity, our data suggests that traditional housing systems may be overlooking a fundamental developmental need. Denying animals the chance to encounter and manage physical challenge during sensitive periods may induce long-lasting deficits in behavioral flexibility and emotional regulation. This risk of a developmental mismatch is heavily supported by the work of Pellis et al. (2023), that demonstrated that a lack of or deficient juvenile play experiences alters the structural development of the prefrontal cortex (Bell et al., 2010; Ham et al., 2024; Omrani et al., 2020; Pellis et al., 2023; Stark et al., 2023) and compromises executive functioning in adulthood (Bijlsma et al., 2022). Integrating elements that mimic natural environmental challenges, such as the wobbly elements used in this study, into standard housing could significantly enhance the ethological validity of rodent models in neuroscience and biomedical research. Ultimately, our quantitative framework provides a methodology to investigate the neurobiological mechanisms underlying risk assessment, motor resilience, and the rewarding value of cognitive-motor challenges during mammalian development. By reframing physical risk-taking not as random, clumsy behavior, but as a highly regulated and essential component of the juvenile play repertoire, this study underscores the importance of environmental unpredictability. Providing young animals with a (safe) space to test the limits of motor skills may be a crucial prerequisite for developing a resilient brain, offering an understanding of how elements in mammalian play functions as a vital preparation for an unpredictable world.

## Notes

### Competing Interest Statement

The authors have declared no competing interest.

### Funding

- Dutch Research Council: OCENW.M.21.083

